# One tagger, many uses: Illustrating the power of ontologies in dictionary-based named entity recognition

**DOI:** 10.1101/067132

**Authors:** Lars Juhl Jensen

**Author notes:** This work was in part funded by the Novo Nordisk Foundation (NNF14CC0001) and the National Institutes of Health (U54 CA189205-01).

## Abstract

Automatic annotation of text is an important complement to manual annotation, because the latter is highly labour intensive. We have developed a fast dictionary-based named entity recognition (NER) system and addressed a wide variety of biomedical problems by applied it to text from many different sources. We have used this tagger both in real-time tools to support curation efforts and in pipelines for populating databases through bulk processing of entire Medline, the open-access subset of PubMed Central, NIH grant abstracts, FDA drug labels, electronic health records, and the Encyclopedia of Life. Despite the simplicity of the approach, it typically achieves 80–90% precision and 70–80% recall. Many of the underlying dictionaries were built from open biomedical ontologies, which further facilitate integration of the text-mining results with evidence from other sources.

## I. Introduction

Named entity recognition (NER) is a fundamental task in biomedical text mining and can benefit greatly from the use of ontologies. This is especially true for dictionary-based NER methods, which with a good ontology at hand can be quickly adapted to a new task without the need for a manually curated corpus for training.

## II. Software Implementation

The core of our NER system is a highly optimized dictionary-based tagging engine implemented in C++ [1]. The core tagger makes use of a custom hashing function to process thousands of PubMed abstracts per second with a single CPU thread. It is furthermore is inherently thread-safe, allowing for perfect scalability in multi-threaded use and is both available as a command-line tool and as a Python module that was generated in part by the Simplified Wrapper and Interface Generator (SWIG). For real-time applications, we have developed a multi-threaded HTTP server that utilizes this Python module to expose the tagger as a RESTful web service, which includes support for the Open Annotation model [2].

The ability to perform real-time tagging enabled us to develop the Reflect [3] and EXTRACT [4] tools, which helps curators identify and extract terms from any web page and was evaluated favourably in the interactive annotation track of BioCreative V [4].

## III. Dictionaries and Application Areas

Software is only one half of a NER system; the other half is the dictionary with all the names that the software matches against the text. When adapting the NER system to a new biomedical application area, the main work required is the construction of a suitable high-quality dictionary and blacklist names not to be tagged. The latter is created through manual inspection of the most frequently occurring dictionary names in a large text corpus.

### A. Molecular Entities

NER and normalization of genes and proteins has been the subject of several BioCreative tasks over the years, most recently in BioCreative III [5]. This was also one of the very first uses of the tagger, which is a key component of the text-mining pipeline in the STRING database [6]. The underlying dictionary of gene/protein names is based on Ensembl [7] and RefSeq [8], which were expanded with additional synonyms from UniProt [9]. An older version of the system achieved F-scores of 91% and 66% for recognition and normalization of yeast and fly genes, respectively [3]. This NER method is also heavily used within the Illuminating the Druggable Genome program to assess how well studied drug targets are based on both publications and NIH RePORTER funding data.

Identification of small-molecule chemical compounds in text was a task in BioCreative V [10]. The tagger is also used for this in the STITCH database [11], which relies on a dictionary constructed from a filtered version of PubChem [12]. STRING and STITCH both employ a statistical cooccurrence as well as natural language processing (NLP) for subsequent extraction of relations between the identified molecular entities from Medline and the open-access subset of PubMed Central. These relations are integrated with evidence from many other sources including experimental data and manually curated pathway databases.

### B. Protein Localization and Expression

The COMPARTMENTS [13] and TISSUES [14] databases take a very similar integrative approach to associate proteins with their subcellular localizations and tissue expression patterns. To this end, we constructed dictionaries based on the cellular component part of Gene Ontology [15] and the Brenda Tissue Ontology [16], respectively. Both ontologies were well populated with synonyms, which were automatically expanded to construct plural and adjective forms. The resulting resources can be used to filter protein networks from STRING to include only proteins from certain subcellular and/or tissue contexts. This is useful, for example, in prediction of host–pathogen interactions.

### C. Diseases and Adverse Drug Reactions

The DISEASES database [17] uses the very same approach to extract disease–gene associations from Medline abstracts. In this case, we run the tagger with a dictionary based on Disease Ontology [18]; this NER approach has been shown to compare favourably with other methods [19].

When treating diseases with drugs, patents may experience adverse drug reactions (ADRs). The SIDER database [20] extracts information on known ADRs from FDA drug labels using an NLP system, which uses the tagger Python module to recognize names from the Unified Medical Language System (UMLS) Metathesaurus for all terms of the Medical Dictionary for Regulatory Activities (MedDRA). In a separate study, we showed that it is also possible to identify ADRs in the clinical narrative text of electronic health records, which required the construction of a separate ADR dictionary in Danish [21]. The latter achieved 89% precision and 75% recall.

### D. Organisms and Environments/Habitats

The applications described so far all fall within molecular biomedicine; however, the tagger has proven equally useful within biodiversity and ecology. Specifically, we have created dictionaries of taxa [1] and environmental descriptors [22] from the NCBI Taxonomy [23] and the Environment Ontology [24], respectively. This achieved 83.9% precision and 72.6% recall for species [1] and 87.8% precision and 77.0% recall for environments [22]. We use these dictionaries with the tagger to extract structured information on habitats of organisms based on their textual descriptions in the Encyclopedia of Life [22].

Most recently we participated in the related 2016 BioNLP shared task on bacterial biotopes, specifically NER of bacteria and biotopes. To this end we implemented rules to refine the match boundaries and normalization of bacterial names and compiled a biotope dictionary by extending the OntoBiotope habitat ontology with additional synonyms from other relevant ontologies [25].

## IV. Conclusions

Despite its simplicity, dictionary-based NER is a powerful approach that in many cases can give comparable performance to more advanced methods, if care is taken when constructing the dictionaries. The dictionary-based approach is particularly attractive in the biomedical domain due to the many ontologies that provide excellent starting points constructing dictionaries.

## References

[1] E. Pafilis, et al., “The SPECIES and ORGANISMS resources for fast and accurate identification of taxonomic names in text,” PLoS One, vol. 8, e65390, 2013.

[2] S. Pyysalo, et al., “Sharing annotations better: RESTful Open Annotation,” Proc. ACL-IJCNLP, pp. 91–96, 2015.

[3] E. Pafilis, et al., “Reflect: augmented browsing for the life scientist,” Nat. Biotechnol., vol. 27, pp. 508–510, 2009.

[4] E. Pafilis, et al., “EXTRACT: Interactive extraction of environment metadata and term suggestion for metagenomic sample annotation,” Proc. BioCreative Challenge Evaluation Workshop, pp. 384–395, 2015.

[5] Z. Lu et al., “The gene normalization task in BioCreative III,” BMC Bioinformatics, vol. 12(S8), S2, 2011.

[6] D. Szklarczyk, et al., “STRING v10: protein-protein interaction networks, integrated over the tree of life,” Nucleic Acids Res., vol. 43, pp. D447–D452, 2015.

[7] F. Cunningham, et al, “Ensembl 2015,” Nucleic Acids Res., vol. 43, pp. D662–D669, 2015.

[8] T. Tatusova, et al., “RefSeq microbial genomes database: new representation and annotation strategy,” Nucleic Acids Res., 42:D553–D559, 2014.

[9] UniProt Consortium, “UniProt: a hub for protein information,” Nucleic Acids Res., vol. 43, pp. D204–D212, 2015.

[10] C.-H. Wei, et al., “Assessing the state of the art in biomedical relation extraction: overview of the BioCreative V chemical-disease relation (CDR) task,” Vol. 2016, baw032, 2016.

[11] M. Kuhn, et al., “STITCH 4: integration of protein-chemical interactions with user data,” Nucleic Acids Res., vol. 42, pp. D401–D407, 2014.

[12] E. Bolton, Y. Wang, P.A. Thiessen, and S.H. Bryant, “PubChem: integrated platform of small molecules and biological activities,” Annu. Rep. Comput. Chem., vol. 4, pp. 217–241, 2008.

[13] J.X. Binder, et al., “COMPARTMENTS:unification and visualization of protein subcellular localization evidence,” Database, vol. 2014, bau012, 2014.

[14] A. Santos, et al., “Comprehensive comparison of large-scale tissue expression datasets,” PeerJ, vol. 3, e1054, 2015.

[15] M. Ashburner, et al., “Gene ontology: tool for the unification of biology. The Gene Ontology Consortium,” Nat. Genet., vol. 25, pp. 25–29, 2000.

[16] A. Chang, et al., “BRENDA in 2015: exciting developments in its 25th year of existence,” Nucleic Acids Res., vol. 43, pp. D439–D446, 2015.

[17] S. Pletscher-Frankild, et al, “DISEASES: text mining and data integration of disease-gene associations,” Methods, vol. 74, pp. 83–89, 2015.

[18] W.A. Kibbe, et al., “Disease Ontology 2015 update: an expanded and updated database of human diseases for linking biomedical knowledge through disease data,” Nucleic Acids Res., vol. 43, pp. D1071–D1078, 2015.

[19] S. ElShal, et al., “A comprehensive comparison of two MEDLINE annotators for disease and gene linkage: sometimes less is more,” Lecture Notes in Computer Science, vol. 9656, pp. 765–778, 2016.

[20] M. Kuhn, I. Letunic, L.J. Jensen, and P. Bork, “The SIDER database of drugs and side effects,” Nucleic Acids Res., vol. 44, pp. D1075–D1079, 2016.

[21] R. Eriksson, et al., “Dictionary construction and identification of possible adverse drug events in Danish clinical narrative text,” J. Am. Med. Inform. Assoc., vol. 20, pp. 947–953, 2013.

[22] E. Pafilis, et al., “ENVIRONMENTS and EOL: identification of Environment Ontology terms in text and the annotation of the Encyclopedia of Life,” Bioinformatics, vol. 31, pp. 1872–1874, 2015.

[23] S. Federhen, “Type material in the NCBI Taxonomy Database,” Nucleic Acids Res., vol. 43, pp. D1086–D1098, 2015.

[24] P. L. Buttigieg, et al., “The environment ontology: contextualising biological and biomedical entities,” J. Biomed. Semant., vol. 4, p. 43, 2013.

[25] H. V. Cook, E. Pafilis, and L. J. Jensen, “A dictionary- and rule-based system for identification of bacteria and habitats in text”, to appear in Proc. BioNLP Shared Task Workshop, 2016.

